# Task-Dependent Performance of GNNs and Pretrained Models in ADMET Prediction

**DOI:** 10.64898/2026.10.26.761652

**Authors:** Martin Stamenov, Mila Gjurovska, Viktorija Vodilovska, Ilinka Ivanoska

**Affiliations:** Faculty of Computer Science and Engineering, Ss Cyril and Methodius University, Skopje, N. Macedonia

**Keywords:** Graph neural networks, ADMET prediction, molecular property prediction, scaffold-split, pretrained, models, drug discovery, benchmarking

## Abstract

Structure-based prediction of ADMET (absorption, distribution, metabolism, excretion, and toxicity) properties promises fast, low-cost screening of drug candidates, yet model selection remains unclear because published comparisons of molecular representations often use unmatched tuning budgets and leakage-prone evaluation protocols, producing performance differences that may not reflect model quality. To address this gap, we benchmark graph neural networks (GNNs), frozen pretrained models (ChemBERTa, MolCLR), and fingerprint baselines across six Therapeutics Data Commons ADMET endpoints under scaffold-split evaluation. To make it really fair we gave every model family a matched 50-trial, validation-selected tuning budget and reported the median over ten seeds; architecture and hyperparameter selection use validation data only, with the held-out test split reserved for final performance estimation. Under this fair comparison protocol, *no model family dominates* and the differences between families fall within the seed-to-seed range on most endpoints. The GNN is clearly best only on Caco-2 permeability (R^2^ ≈ 0.48), and a controlled ablation shows this advantage comes entirely from auxiliary physicochemical descriptors rather than the graph representation; Morgan fingerprints are competitive on hERG (AUC ≈ 0.77) and give the best median on both clearance endpoints, while MolCLR is best on Tox21. For hepatocyte clearance and half-life, all models yield near-zero or negative R^2^. No single hyperparameter-optimization strategy, among seven evaluated, consistently outperforms random search at this budget, and we show that selecting the optimizer using test-set performance or reporting only single best-trial runs can substantially inflate a favored model’s apparent advantage. Reliable ADMET modeling therefore requires endpoint-specific model selection together with fair, leakage-controlled, seed-averaged evaluation, offering more trustworthy decision support for early-stage drug candidate prioritization.

## I. Introduction

Predicting molecular ADMET (Absorption, Distribution, Metabolism, Excretion, and Toxicity) properties is a central task in computational drug discovery, enabling early identification of promising candidates and flagging safety liabilities before costly synthesis and *in vivo* testing [1]. Graph

Neural Networks (GNNs) have become prominent in this domain by treating molecules as graphs in which atoms serve as nodes and bonds as edges, enabling task-specific learning through message passing that encodes both local chemical motifs and global topological patterns inaccessible to fixed-length descriptors [2]–[4].

Beyond GNNs, pretrained encoders such as ChemBERTa [5] and MolCLR [6] leverage self-supervised learning on large corpora, while Morgan (ECFP) fingerprints [7] paired with MLP predictors remain competitive baselines [8], [9]. Although these approaches differ in architecture and encoding strategy, all model families evaluated here rely exclusively on molecular structure as input, representing the same underlying 2D molecular graph or SMILES string [10] through different feature representations. Across these paradigms, strong predictive performance has been reported, often achieving state-of-the-art results on individual tasks [11].

Despite this progress, model selection in practice remains unclear. Different models perform inconsistently across tasks, and most published studies optimize performance for individual architectures or compare methods under different splits and experimental conditions, making direct comparison unreliable [3], [12]. Crucially, many works rely on random data splits, which allow structurally similar molecules to appear in both training and test sets, producing overly optimistic estimates that do not reflect the real-world scenario of predicting properties for new chemical scaffolds. As a result, practitioners still lack consistent guidance on model selection, endpoint-specific reliability, and the conditions under which molecular structure alone provides sufficient predictive information [11], [12].

Scaffold-based splitting [13] partitions compounds by core ring structures and serves as a realistic evaluation protocol reflecting prospective drug discovery, where models must generalize to novel chemical series. In this work, we investigate how predictive performance depends on the underlying task and evaluation protocol, with a focus on identifying when molecular structure alone provides sufficient signal. This issue is particularly important for pharmacokinetic endpoints such as clearance and half-life, where inaccurate predictions can affect compound prioritization. Models that appear accurate under weak evaluation protocols may provide misleading support for early-stage screening decisions.

Benchmarking should therefore identify not only which model performs best, but also which endpoints are predictable from molecular structure under realistic scaffold-shift conditions, motivating endpoint-specific model selection according to the biological and experimental characteristics of each endpoint.

The contributions of this paper are: i) a matched-budget, ten-seed benchmark of GNN, pretrained, and fingerprint models on six scaffold-split TDC ADMET endpoints; ii) an endpoint-level analysis of where molecular structure alone carries usable signal; and iii) a separate comparison of seven HPO strategies against Random Search, kept apart from the model-family comparison.

The remainder of this paper is organized as follows. Section II reviews related work. Section III describes materials and methods. Section IV presents experimental results. Section V discusses implications and limitations. Section VI concludes.

## II. Related Work

The TDC ADMET benchmark [11] has become the standard evaluation platform for molecular property prediction, and its leaderboards [14] show that comparatively simple methods remain highly competitive. Three paradigms dominate. *Fingerprint and descriptor methods* pair Morgan or RDKit2D features with MLPs, random forests, or SVMs (MapLight [8], RDKit2D+MLP [9]), and ensembles of such models are also strong [15]. *Graph neural networks* learn task-specific representations end-to-end, including GCN [16], GIN [17], the bond-level D-MPNN [3], and the attention-based AttentiveFP [4]. *Pretrained encoders* use self-supervised objectives on large unlabeled corpora, either attribute masking [18] or contrastive learning [19], followed by optional multi-task training and task-specific fine-tuning. Examples include the contrastive graph model MolCLR [6] (atom masking, bond deletion, subgraph removal), the SMILES transformers ChemBERTa [5] (RoBERTa pretrained on 77M ZINC-15 [20] molecules) and MolBERT [21], and DeepMol [22], MolE [23], and the compact MiniMol [24] (∼ 10M parameters).

Moreover, transformer-based models frequently show severe generalization drops under scaffold splitting, with strong validation scores that collapse toward chance-level performance on held-out scaffolds [3], [5]. Wu et al. [12] introduced MoleculeNet with scaffold-splitting to reflect prospective drug discovery generalization; we revisit this collapse in Section IV.

HPO is a key performance driver in molecular ML, as well, where small, noisy datasets amplify sensitivity to hyperparameter configuration. Bergstra and Bengio [25] established Random Search as a strong baseline, showing it outperforms grid search when only a subset of hyperparameters is influential. Bayesian methods such as TPE [26], implemented in Optuna [27], improve sample efficiency by modeling the hyperparameter-performance relationship. Population-based metaheuristics, including PSO [28], GA [29], and ABC [30], provide broader exploration but may require larger budgets to converge. SA [31] offers a middle ground by accepting suboptimal moves early to escape local optima. Despite their widespread use in combinatorial optimization, systematic comparisons of these HPO strategies for molecular property prediction remain scarce, particularly under scaffold-split protocols, motivating our evaluation within a unified ADMET benchmark.

In all, ADMET prediction remains challenging because many endpoints exhibit distribution shifts under scaffold splitting, limited dataset sizes, and class imbalance. These factors can substantially affect model rankings and generalization estimates, motivating the use of multi-seed evaluation and confidence-interval-based reporting throughout this study.

## III. Materials and Methods

### A. Framework Overview

To compare model families under realistic generalization conditions, we use a benchmarking framework (Fig. 1) that applies the same scaffold partitions, datasets, and metrics across the GNN and the frozen-pretrained/fingerprint pipelines. The framework coordinates dataset loading, scaffold-split partitioning, training, metric computation, multi-seed analysis, and HPO comparison against Random Search. Fixing splits and metrics across models reduces experimental confounds and enables direct comparison between model families [12].

**Fig. 1.**
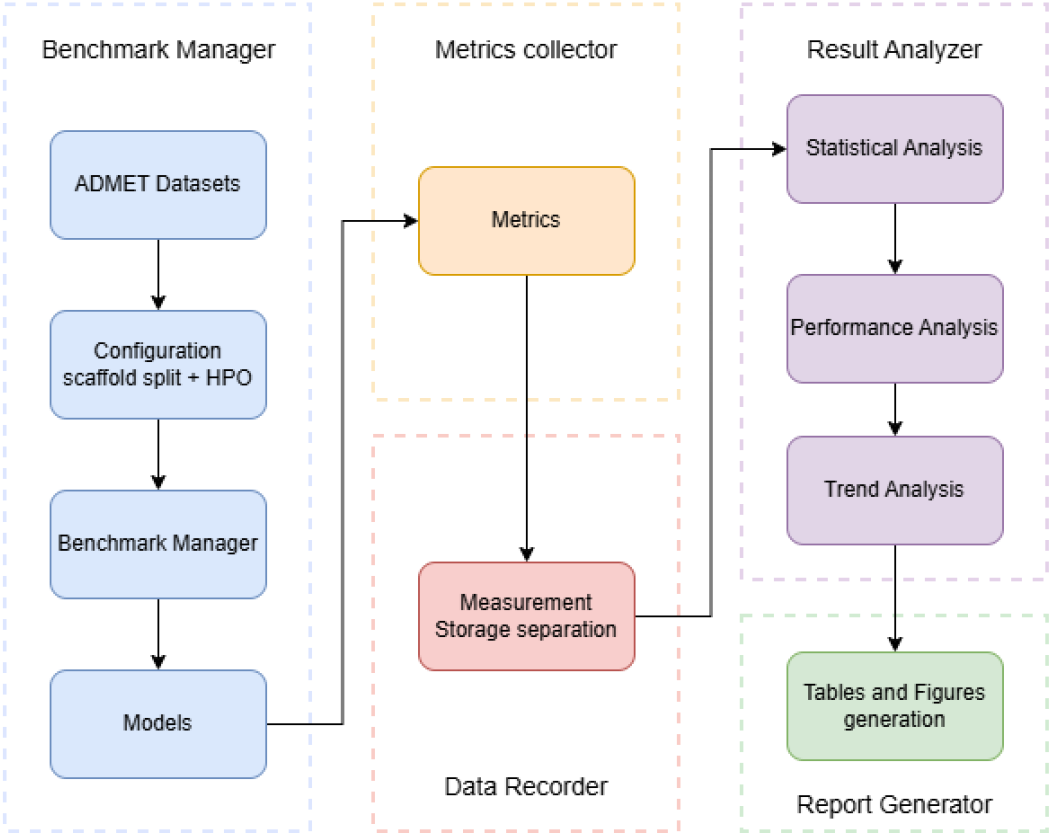
Benchmarking framework: orchestration, model execution, and evaluation layers.

### B. Datasets and Evaluation Protocol

We evaluate all models on six molecular property prediction tasks drawn from the TDC ADMET benchmark [11], covering four pharmacokinetic (ADME) regression tasks and two toxicity classification tasks. Of the 11,805 catalogued compounds, 10,627 were retained after removing molecules with invalid SMILES or failed graph conversion (Table I). This protocol reflects practical drug discovery scenarios, where models must generalize to novel chemical series.

**TABLE I.** Dataset statistics and evaluation metrics (TDC ADMET). “TDC” shows compounds catalogued and “Used” shows those retained after filtering. RMSE is the primary per-dataset regression metric, the scale-invariant R***2*** is additionally used for cross-model comparison in Table II.

| Dataset | Type | TDC | Used | Split | Metric |
| --- | --- | --- | --- | --- | --- |
| Caco2_Wang | Regr. | 910 | 819 | Scaffold | RMSE |
| Half_Life | Regr. | 667 | 601 | Scaffold | RMSE |
| Clear_Hep. | Regr. | 1,213 | 1,092 | Scaffold | RMSE |
| Clear_Mic. | Regr. | 1,102 | 992 | Scaffold | RMSE |
| Tox21 (NR-AR) | Class. | 7,258 | 6,533 | Scaffold | AUC |
| hERG | Class. | 655 | 590 | Scaffold | AUC |
| Total |  | 11,805 | 10,627 |  |  |

The ADME regression tasks are evaluated on the following datasets: i) Caco-2 Permeability (Caco2_Wang) (contains 910 compounds with permeability values reported in log(cm/s) units and is commonly used as an *in vitro* surrogate of intestinal absorption [32]); ii) Half-Life Prediction (Half_Life_Obach) (contains 667 compounds with plasma half-life measurements. Target values are log-transformed prior to standardization to reduce skewness.); iii) Hepatocyte Clearance (Clearance_Hepatocyte_AZ) (AstraZeneca dataset containing 1,213 compounds with intrinsic clearance measurements obtained from human hepatocyte assays [11].); and iv) Microsomal Clearance (Clearance_Microsome_AZ) (AstraZeneca microsomal clearance dataset containing 1,102 compounds with human liver microsome measurements. Targets are log-transformed and standardized using training-set statistics.).

The Toxicity Classification Tasks are evaluated on: i) Tox21 Nuclear Receptor: Androgen Receptor (Tox21 NR-AR) (contains 7,258 compounds with binary androgen receptor activity labels and exhibits substantial class imbalance, with approximately 4.2% positive samples in the full corpus (rising to ≈4.9% on the scaffold-split test set, see Section IV)); and ii) hERG Cardiotoxicity (hERG) (contains 655 compounds with blocker/non-blocker labels and is widely used for cardiotoxicity assessment [33].)

TDC scaffold-split function is used, which partitions compounds by Bemis-Murcko scaffolds [13] so that molecules with structurally distinct core rings appear in different folds. This approximates prospective deployment conditions, where a model must generalize to structurally distinct compound families not seen during training. We further subdivide the training partition into training and validation using a fixed random seed (42) at the molecule level, so no molecule occurs in both; 10% of unique molecules form the validation set for early stopping and model selection. The final split proportions are approximately 70/8/22 (train/validation/test) across all datasets. During HPO, the test split remains *fully held out*, model selection is performed solely on the validation metric. Final performance is reported once on the test split after HPO is complete, preventing any implicit information leakage from repeated test-set evaluations.

For Half-Life, Hepatocyte, and Microsomal Clearance, targets are log-transformed using *y*^*′*^ = log(max(*y, ϵ*)), *ϵ* = 10^−3^, followed by training-set z-score normalization. Caco-2 targets require only normalization, while classification labels remain binary.

### C. Model Families

We evaluate three distinct model paradigms under the same scaffold splits and evaluation metrics, allowing us to assess whether task-specific learning provides advantages over frozen pretrained representations and fixed fingerprint baselines.

Firstly, we employ graph neural networks trained end-to-end on each task. The GNN backbone is based on stacked GraphConv layers [34] with batch normalization, ReLU activation, and additive residual connections, selected through a preliminary architecture comparison across eight GNN variants (Section IV-D). Each molecule is represented as a molecular graph constructed from SMILES using RDKit. Atom features include standard atom-level chemical descriptors (8 features); no bond features are used. The GNN encoder produces node embeddings aggregated into a graph-level representation using concatenated global mean and max pooling. Before the prediction head, this pooled representation is concatenated with a small vector of structure-derived physicochemical descriptors computed with RDKit (15 for the ADME/toxicity tasks, e.g. molecular weight, cLogP, TPSA, hydrogen-bond donors/acceptors, and Lipinski flags; a 7-descriptor subset for Caco-2). These descriptors carry no biological or assay information and are provided only to the GNN, not to the pre-trained or fingerprint baselines. The combined representation is passed to an MLP prediction head with batch normalization and optional dropout; for the small Caco-2 dataset the GNN additionally applies dropout (0.5) and Gaussian target-label noise (*σ* = 0.05). GNN models undergo full hyperparameter optimization (Section III-D).

Furthermore, we evaluate two pretrained molecular encoders representing the two major pretrained paradigms: Chem-BERTa [5] (SMILES-based transformer) and MolCLR [6] (graph-based contrastive model), as representative rather than exhaustive examples. For pretrained encoders, we use a frozen encoder and trainable MLP head setting: embeddings are extracted once per molecule, and an MLP is trained on the train split with early stopping based on the validation split. This frozen-encoder protocol isolates the quality of pretrained representations from downstream optimization effects.

Two fingerprint-based approaches are evaluated: (i) Morgan fingerprints (ECFP) [7] with an MLP predictor, and (ii) MolE-style learned fingerprints (MolE-FP) with an MLP predictor. The MolE-FP baseline is implemented as Morgan fingerprints followed by a learned projection inspired by the MolE approach [23], rather than the original pretrained MolE encoder. All baselines use the same dataset splits and evaluation metrics as the GNN experiments.

### D. Model Training and Configuration

a. *GNN Training:* All GNN models were trained using the Adam optimizer with a batch size of 32. During HPO, models were trained for a maximum of 50 epochs per trial with early stopping (patience = 12). Regression models used mean squared error (MSE) loss, while classification models used binary cross-entropy (BCE) loss with logits. Gradient clipping was applied with a maximum norm of 1.0. The hyperparameter search space is defined in Supplementary Materials (Table S2).
b. *Optimization algorithms:* We evaluate seven HPO strategies: Random Search [25], Particle Swarm Optimization (PSO) [28], Artificial Bee Colony (ABC) [30], Genetic Algorithm (GA) [29], Simulated Annealing (SA) [31], Hill Climbing (HC) [35], and Tree-structured Parzen Estimator (TPE) [27]. Population-based methods use population size 16. SA uses *T*_0_ = 1.0 with exponential cooling (*α* = 0.99), while TPE uses Optuna defaults with median pruning. Search spaces are provided in Supplementary Table S2. Each algorithm is allocated 50 trials per dataset, giving 2,100 GNN training runs from the optimizer comparison alone (seven optimizers *×* six datasets *×* 50 trials). In addition, the study includes a preliminary architecture-selection screen (reported in the Supplementary Material) and 60 multi-seed validation runs for the final selected models. We keep two questions separate. For the *fair model-family comparison* (Table II), the GNN is tuned with a single predefined optimizer, Random Search, whose configuration is selected on validation and evaluated once on test, matching the 50-trial validation-based budget given to every baseline; the test set is never used to choose the optimizer. The seven-optimizer study is a *separate* analysis of whether optimizer choice affects the GNN, and its outcomes do not feed the model-family comparison.

**TABLE II.** Fair model-family comparison on six scaffold-split ADMET endpoints: median test performance over ten seeds of each family’s validation-selected configuration (R^2^ for regression, AUC-ROC for classification). Every family receives a matched 50-trial, validation-based tuning budget (GNN: Random Search over architecture and training hyperparameters; pretrained and fingerprint models: search over the MLP head on frozen features), and no test information is used for selection. Bold marks the highest median, but leading families typically differ by less than the across-seed range (Section IV-C). Negative R^2^ means worse than predicting the test-set mean; medians are reported because some configurations diverge on a minority of seeds.

| Task | Metric | GNN | ChemBERTa | Morgan-FP | MolE-FP | MolCLR |
| --- | --- | --- | --- | --- | --- | --- |
| *Group 1: Structure-driven tasks* |  |  |  |  |  |  |
| hERG | AUC ↑ | 0.754 | 0.747 | **0.767** | 0.735 | 0.657 |
| Caco-2 | $R^2$ ↑ | **0.482** | 0.413 | 0.322 | 0.127 | 0.232 |
| *Group 2: Partially structure-driven tasks* |  |  |  |  |  |  |
| Tox21 (NR-AR) | AUC ↑ | 0.718 | 0.714 | 0.706 | 0.698 | **0.722** |
| Clearance Mic. | $R^2$ ↑ | 0.093 | −0.010 | **0.169** | 0.056 | 0.125 |
| *Group 3: Tasks weakly learnable from structure alone* |  |  |  |  |  |  |
| Half-Life | $R^2$ ↑ | −**0.010** | −1.918 | −6.341 | −3.776 | −0.672 |
| Clearance Hep. | $R^2$ ↑ | −0.373 | −0.224 | −**0.017** | −0.106 | −0.113 |

### E. Experimental Setup

Table I summarizes the dataset characteristics across all six tasks. Model performance is assessed using RMSE, R^2^, and MAE for ADME regression tasks, and AUC-ROC and F1-score for toxicity classification tasks. AUC-ROC is the primary ranking metric for classification and is the classification metric used in the paired optimizer comparison (Supplementary Table S4); F1 is reported in addition because it is more sensitive to the minority class under strong class imbalance.

To quantify robustness, we retrain the best HPO-selected configuration per dataset across ten seeds [42, 123, 456, 789, 1011, 2022, 3033, 7, 99, 2718]. For the across-seed results (Section IV-C and Table III), we use the median and full min–max range across seeds because the selected configuration diverges on a minority of seeds for some endpoints, making the mean unrepresentative. These medians provide a more conservative estimate than the optimistic best-of-50-trials result. Secondary imbalance-aware toxicity metrics (PR-AUC, F1, MCC) are reported as mean *±* standard deviation to characterize variability.

**TABLE III.** Across-seed instability of the GNN’s validation-selected configuration (GraphConv backbone) on the two clearance endpoints, as median [min–max] RMSE over ten seeds. The configuration diverges on a minority of seeds (strongly negative R^2^); the Hepatocyte maximum is roughly 20 times its minimum.

| Dataset | Metric | Median [min-max] |
| --- | --- | --- |
| Clear_Hep. | RMSE | 56.22 [46.53–977.95] |
| Clear_Mic. | RMSE | 41.03 [38.48–55.37] |

We compare each optimizer against the Random Search baseline using paired per-endpoint differences across the six datasets rather than null-hypothesis significance tests. For every endpoint we take the difference in that endpoint’s primary test metric (R^2^ for regression, AUC-ROC for classification), oriented so that positive values favor the optimizer. Because these metrics lie on incomparable scales and the regression R^2^ is occasionally pathologically negative, a raw mean difference is not a sound summary; we therefore report (i) the win/tie/loss count with an exact two-sided sign test on wins-versus-losses (ties dropped), (ii) a scale-free, direction-normalized paired effect, each endpoint’s difference divided by the spread of all seven optimizers on that endpoint, then averaged over the six endpoints, and (iii) two sensitivity variants that recompute the mean paired difference with R^2^ clipped to [−1, 1] and floored at 0 (AUC untouched). All 95% confidence intervals are 10,000-resample percentile bootstraps over the six paired differences. The paired unit is the dataset (*n* = 6), and all comparisons use final held-out test performance obtained after HPO model selection on the validation split. With only six paired observations, null-hypothesis significance tests are underpowered, so we read these measures descriptively, summarizing the magnitude and the consistency of any improvement over Random Search (Supplementary Table S4).

Experiments were implemented in Python using PyTorch, PyTorch Geometric, RDKit, PyTDC, NiaPy, Optuna, and Transformers. Training was performed on a workstation equipped with an Intel Core i7-8700K CPU, 16 GB RAM, and an NVIDIA RTX 3060 GPU.

## IV. Results

### A. Task-Level Performance on ADMET Endpoints

Table II presents the performance of all model families across the six ADMET tasks. Fig. 2 provides a visual summary of relative model rankings. We observe substantial variability in predictive performance across tasks, which we organize into three groups based on the degree to which molecular structure supports prediction.

**Fig. 2.**
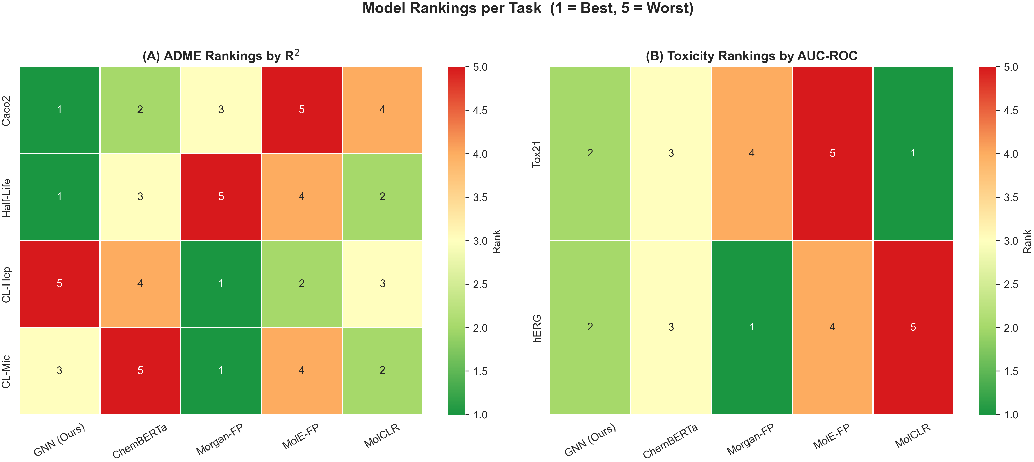
Task-dependent performance of different model families across ADMET endpoints. Green indicates better performance within each task.

a. *Group 1: Structure-driven tasks*.: On hERG and Caco-2 the models reach their highest scores, but no single family is clearly best. On hERG the leading families lie within 0.03 AUC of one another (Morgan-FP highest at 0.77, with the GNN and ChemBERTa at 0.75), a spread smaller than the across-seed range, so we do not read any as a significant winner. On Caco-2 the GNN attains the highest median R^2^ (0.48), though this edge is descriptor-driven (Section IV-B). These endpoints have relatively strong structure-activity relationships: hERG blockade is associated with recurring pharmacophoric features [36], and intestinal permeability tracks molecular size, lipophilicity, polar surface area, and hydrogen-bond capacity [37], [38].
b. *Group 2: Partially structure-driven tasks*.: On Tox21 and Microsomal clearance predictive power is moderate and, again, closely matched. On Tox21 all families lie between AUC 0.69 and 0.73 (MolCLR highest at 0.722, GNN 0.718); on Microsomal clearance Morgan-FP gives the highest median R^2^ (0.17), ahead of MolCLR (0.13) and the GNN (0.09). Structure is partially informative here, but no model family dominates.
c. *Group 3: Tasks weakly learnable from structure alone*.: On Half-Life and Hepatocyte clearance all families produce near-zero or negative R^2^. The best Half-Life median R^2^ is only ≈ 0 (−0.01, GNN), and every other family is negative, strikingly so for the fingerprint baselines, whose validation-selected heads overfit these tiny scaffold-split training sets and collapse to large negative test R^2^ (Morgan-FP −6.3, MolE-FP −3.8). On Hepatocyte clearance the best median is Morgan-FP (−0.02); the GNN’s validation-selected configuration is unstable on this endpoint and posts a strongly negative median R^2^ (− 0.37), with individual seeds reaching strongly negative raw R^2^ values (Table III). These results indicate that, for the structure-only representations evaluated here under scaffold splitting, molecular structure alone does not support robust prediction of these pharmacokinetic endpoints, which depend on enzyme kinetics, transporter activity, protein binding, and individual patient variability [39]. The similarly weak performance across simple and complex models points to a limitation in the information content of structure-only representations rather than the choice of learning algorithm.

### B. Model Performance Across Tasks

a. *The GNN’s one advantage is descriptor-driven*.: The GNN alone receives auxiliary physicochemical descriptors that the frozen baselines do not: a compact set of 15 RDKit-computed features (7 for Caco-2), rather than the 200-dimensional RDKit2D block used by the RDKit2D+MLP baseline. A controlled ablation (each endpoint’s validation-selected configuration retrained with and without the descriptors, median over ten seeds) shows this input is decisive precisely on the one endpoint where the GNN leads: on Caco-2 the median test R^2^ falls from 0.45 with descriptors to −0.51 without them (Δ ≈ 0.96–0.97 depending on rounding of intermediate values), so the GNN’s Caco-2 advantage is attributable to the descriptors rather than to graph message passing. On the two toxicity endpoints the descriptors add essentially nothing (hERG ≈ Δ +0.00, Tox21 ≈ Δ +0.01): the graph model alone (hERG ≈ 0.76) already performs at the level of the fingerprint and pretrained baselines. On the weakly-learnable clearance and half-life endpoints the effect is small or mixed (Half-Life Δ ≈ +0.02; Hepatocyte clearance Δ ≈ −0.31, both models near or below zero). Once tuning and inputs are matched, graph message passing itself confers no consistent advantage.
b. *Diagnostic for weakly learnable endpoints*.: Frozen representations degrade less than end-to-end GNN training on Hepatocyte clearance (tuned ChemBERTa R^2^ = −0.22 vs. −0.37), so they are best viewed as regularized baselines rather than evidence that pretrained models are superior. In practice, if a task-specific GNN yields negative R^2^ or near-random AUC on a scaffold-split test set, the endpoint may not be predictable from structure alone, and frozen encoders provide a useful sanity check.

### C. Effect of Evaluation Protocol

*Model-family differences are small under scaffold-split evaluation*.: Across ten seeds, the GNN’s validation-selected configuration is relatively stable on hERG (median AUC 0.754, range 0.704–0.809) and Tox21 (0.718, 0.695–0.749), and these ranges exceed the gaps between the leading families in Table II. It is likewise stable on Caco-2 and Half-Life, but on Microsomal and Hepatocyte clearance it diverges on a minority of seeds, with individual seeds reaching strongly negative R^2^ (Table III). We therefore report medians, which are robust to these runs; the wide clearance ranges are consistent with our central finding that these pharmacokinetic endpoints do not support robust structure-only prediction. Imbalance-aware toxicity metrics (PR-AUC, MCC, F1) are summarized separately as mean *±* standard deviation in the Supplementary Material.

a. *Evaluation protocol influences model rankings*.: Scaffold-based splitting creates a structural covariate shift where the model must generalize not merely to unseen molecules, but to unseen chemical families. Under such conditions, validation performance becomes a noisier proxy for true generalization, because a configuration that fits the specific scaffolds in the validation set may not transfer to the test scaffolds. On Tox21, the pretrained encoders reach high validation AUC that does not carry over to the scaffold-split test set: ChemBERTa drops from validation AUC ≈ 0.92 to a test median of 0.714, and MolCLR from ≈ 0.90 to 0.722 (Table II). These results demonstrate that evaluation protocol substantially influences model rankings and perceived performance. Partial fine-tuning does not change this picture: unfreezing ChemBERTa’s last two transformer layers (evaluated on the same canonical split as every other model) yields test performance comparable to the frozen-feature protocol, within a few points on the classification endpoints (hERG AUC 0.770 vs. 0.747 frozen; Tox21 0.735 vs. 0.714), positive only on Caco-2 (R^2^ = 0.39), and near-zero or negative on the harder regression endpoints (Half-Life −0.08, Hepatocyte clearance −0.25). Fine-tuning neither rescues the difficult pharmacokinetic tasks nor overturns the frozen-feature ranking, so the frozen protocol used for the pretrained models does not understate their performance here.

### D. Model Architecture Selection

A preliminary architecture screening was performed across eight graph neural network variants to select a backbone for the main benchmark, with each architecture evaluated at its best of five validation-selected configurations under the same scaffold-split pipeline. No single architecture dominated: ranked by *validation* performance (the only leak-free selection signal), the average regression rank spans a narrow band (3.0–6.0 across eight backbones) and per-dataset validation differences are small (e.g., Caco-2 validation R^2^ = 0.51– 0.63, Tox21 AUC = 0.78–0.85, hERG AUC = 0.88–0.92), while on the weakly-learnable clearance and half-life endpoints every architecture is unstable (Supplementary Tables S1 and S3). This validation ordering is a poor predictor of test performance, since several validation-competitive backbones post negative mean test R^2^, reinforcing that predictability is endpoint-driven rather than architecture-driven. Because no backbone is meaningfully better on validation, we adopted GraphConv as a single representative message-passing backbone; it is mid-ranked on validation (regression validation rank 4.25 of 8) and is not a test-maximizing choice (its mean test R^2^ is in fact among the lowest), and because selection used validation metrics only, the main-benchmark test evaluation remains unbiased. Full per-architecture results for both the classification and regression endpoints (all eight backbones, evaluated under the same scaffold-split pipeline) are provided in the supplementary material (Tables S1 and S3, Figs. S3 and S4).

### E. Effect of Hyperparameter Optimization

Each optimizer’s configuration is selected on validation and evaluated once on the held-out test set, then compared with Random Search by paired difference across the six datasets (Supplementary Table S4; positive values favor the optimizer). No metaheuristic consistently outperforms Random Search: no optimizer is significant under the paired sign test (the most favorable, ABC, wins on four of six endpoints with one tie, *p* = 0.375), and every scale-free and scale-clipped 95% bootstrap interval spans zero. Only the heavily compressed floor-0 variant separates two optimizers, ABC at a boundary (+0.032, 95% CI [+0.000, +0.091]) and SA slightly below Random Search (− 0.018, 95% CI [− 0.031, − 0.006]); neither indicates a consistent winner. We deliberately do not name a per-dataset “best” optimizer, since such winners are unstable across seeds and must not be chosen with test data. Two factors plausibly explain Random Search’s competitiveness: when only a subset of hyperparameters matters (“effective dimensionality” [25]), random sampling reaches near-optimal configurations at a rate adaptive methods do not reliably exceed in 50 trials; and on noisy scaffold-split validation sets, adaptive methods risk overfitting the validation fold, whereas Random Search does not condition on earlier evaluations. Fig. 3 compares all algorithms on test R^2^ (regression) and AUC-ROC (classification).

**Fig. 3.**
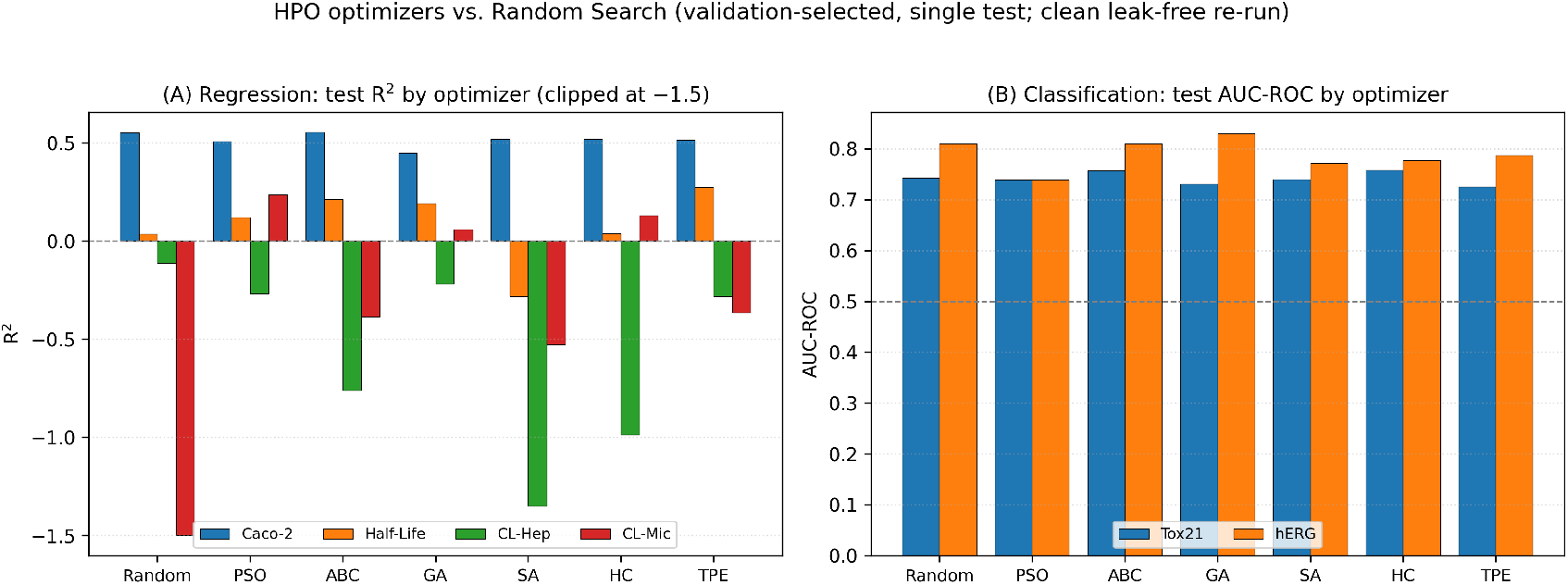
HPO optimizer comparison (clean, leak-free re-run): each optimizer’s *validation-selected* configuration evaluated once on test. (A) Regression test R^***2***^ per optimizer across the four ADME endpoints (clipped at **−1.5** for readability; dashed line at ***R***^***2***^ **= 0**); (B) classification test AUC-ROC across the two toxicity endpoints (dashed line at the **0.5** random baseline). No optimizer is consistently ahead of Random Search (Supplementary Table S4 for the statistics).

The GNN’s validation-selected configuration reaches ROC-AUC *>* 0.7 on both classification tasks. On hERG it achieves F1 = 0.79 (PR-AUC 0.89, MCC 0.37), with balanced performance across both classes. On Tox21 the same configuration reaches F1 = 0.44 and MCC 0.49 despite high overall accuracy, reflecting the severe class imbalance (≈5% positive rate on the scaffold-split test set). Per-class confusion matrices are shown in Supplementary Materials (Fig. S1). Because ROC-AUC can be misleading under severe class imbalance, we treat PR-AUC as a co-primary metric for Tox21 and additionally report MCC [40]; across-seed stability is shown in Fig. S2.

## V. Discussion

The six endpoints span a wide range of learnability, confirming that ADMET modeling should not be treated as one homogeneous problem: each endpoint combines different physicochemical, biological, and experimental factors, and these determine how much signal molecular structure alone can supply.

On Caco-2 and hERG, topology and local chemical environment carry useful signal, and atom-level message passing (our GNN uses atom features and connectivity but no bond features), SMILES transformers, and simple fingerprints capture it comparably; the GNN’s Caco-2 edge is descriptor-driven (Section IV-B), not a message-passing effect.

Half-Life and Hepatocyte clearance instead depend on factors absent from structure, including enzyme-mediated metabolism, transporter interactions, tissue distribution, and plasma protein binding [41]–[43]. The uniformly poor results are therefore better explained by the information content of structure-only inputs than by architecture: added model complexity did not compensate for missing biological context.

The two toxicity tasks differ in class balance and mechanism, which again argues for endpoint-specific evaluation. Severe imbalance on Tox21 NR-AR (≈4.2% positives overall, ≈4.9% in the test split) biases standard objectives toward the majority class [44], so high ROC-AUC should be read cautiously and we treat PR-AUC as co-primary. Future work should add sensitivity, specificity, and calibration error, especially for safety endpoints where false negatives are costly.

Scaffold splitting removes close structural analogues from the test set, so the modest scores observed reflect the genuine difficulty of generalizing beyond familiar chemotypes, a central challenge in drug discovery. Under this protocol no representation is uniformly optimal: frozen pretrained encoders perform according to the alignment between their pretraining objective and the endpoint, and fingerprint models stay competitive when data are small or the target tracks known physicochemical properties. These results favor endpoint-specific model selection over a single architecture.

Because validation performance is noisy on small ADMET datasets, expensive search methods should be justified by consistent gains across independent runs; simple strategies such as Random Search may suffice (Section IV-E). ADMET predictors are therefore best treated as endpoint-specific decision-support tools: structure-only models may support hERG and Caco-2 screening, whereas clearance and half-life predictions require additional biological context.

This study has several limitations. It covers six TDC endpoints, which are important but not exhaustive; the datasets are small and, for toxicity, class-imbalanced, and scaffold splitting, though more realistic than random splitting, remains retrospective rather than prospective validation, so variance across splits is high and conclusions apply only within the evaluated datasets, model families, and protocol. All model families use structure-only inputs (no assay conditions or enzyme, transporter, and protein-binding data), and the GNN alone receives structure-derived descriptors, so the comparison is not strictly input-matched. The pretrained encoders (Chem-BERTa, MolCLR) are representative rather than exhaustive and were evaluated mainly frozen; a partial fine-tuning check (Section IV-C) performed comparably, but a broader study remains future work. Finally, the HPO comparison is limited to 50 trials, and a larger budget could alter metaheuristic rankings; future extensions include multimodal representations and uncertainty-aware evaluation.

## VI. Conclusion

We benchmarked graph neural networks, frozen pretrained molecular representations, and fingerprint baselines across six TDC ADMET endpoints under scaffold-split evaluation, with a matched 50-trial validation-based tuning budget per family and ten-seed evaluation. Performance varied substantially across endpoints, moderate on hERG and Caco-2, near-zero on Half-Life and Hepatocyte Clearance, but *no single model family dominated*: once tuning, seeds, and the evaluation split are matched, the leading families are separated by less than the across-seed range on most endpoints, and simple Morgan fingerprints are competitive throughout. The GNN’s lone edge (Caco-2) is descriptor-driven.

Scaffold-split evaluation is critical for drawing realistic conclusions from ADMET benchmarks. Random splits can produce overly optimistic estimates by allowing structurally similar molecules into both training and test sets, whereas scaffold splitting provides a stricter assessment of generalization to unseen scaffolds. Under scaffold-split evaluation, model-family differences are small and model rankings are endpoint-dependent. We further show that no HPO algorithm consistently outperforms Random Search at a 50-trial budget, with paired 95% confidence intervals that span zero, reinforcing that evaluation protocol strongly influences the apparent benefit of both model architecture and optimization methods.

Future work should move beyond structure-only prediction by incorporating protein targets, assay conditions, experimental metadata, and pharmacokinetic context, and should investigate uncertainty-aware evaluation protocols that better reflect prospective drug discovery. The three learnability levels (structure-driven: hERG, Caco-2; moderately: Tox21 NR-AR, Microsomal Clearance, weakly: Half-Life, Hepatocyte Clearance) are descriptive summaries, not universal thresholds. Two common shortcuts, test-set optimizer selection and single best-trial reporting, can substantially inflate apparent advantages, and controlling for both was necessary to reach the conclusions above. Reliable ADMET modeling therefore requires endpoint-specific model selection together with fair, leakage-controlled, seed-averaged scaffold evaluation, supporting a cautious use of molecular AI as decision support in early-stage drug discovery rather than as a universally reliable predictor.

## Data and Code Availability

All datasets used in this study are publicly available through the Therapeutics Data Commons (TDC). The source code, trained model configurations, hyperparameter-optimization logs, and experimental results are available at: https://github.com/NitramVonemats/ADMET-Benchmark

Multi-seed validation used the fixed seeds [42, 123, 456, 789, 1011, 2022, 3033, 7, 99, 2718], and each HPO algorithm was run for 50 trials per dataset. Software dependencies and exact versions are documented in the repository.

## Ethical Compliance

This study uses only publicly available benchmark datasets from the Therapeutics Data Commons and does not involve human subjects, identifiable patient data, animal experiments, or newly collected clinical data. Therefore, institutional review board approval and informed consent were not required.

## Acknowledgment

This work was supported by the Faculty of Computer Science and Engineering, Ss Cyril and Methodius University, Skopje, N. Macedonia.

---

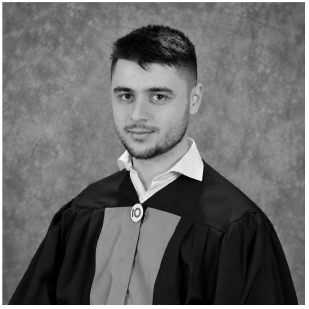

**Martin Stamenov** received the B.Sc. degree in Computer Science from the Faculty of Computer Science and Engineering, Ss. Cyril and Methodius University in Skopje, North Macedonia. He has professional experience in AI engineering, focusing on the development of intelligent systems. His research interests include machine learning, deep learning, graph neural networks, and large language models.

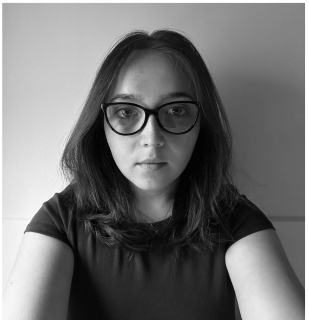

**Mila Gjurovska** received the B.Sc. degree in Software Engineering and Information Systems, from the Faculty of Computer Science and Engineering, Ss. Cyril and Methodius University in Skopje, North Macedonia. She is currently pursuing the M.Sc. degree in Bioinformatics for Health Sciences at the Pompeu Fabra University and the University of Barcelona, Barcelona, Spain. Her research interests include bioinformatics, deep learning, and computational approaches to biomedical data analysis.

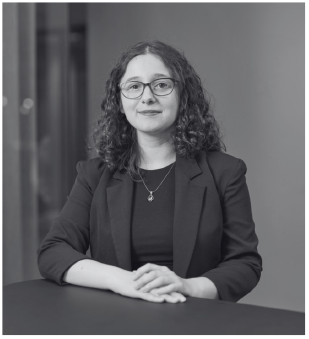

**Viktorija Vodilovska** received the B.Sc. degree in Software Engineering and Information Systems from Ss. Cyril and Methodius University, Skopje, North Macedonia, and the M.Sc. degree in Bioinformatics and Biocomplexity from Utrecht University, The Netherlands. Her research interests include applied machine learning for drug discovery, graph neural networks and computational biology.

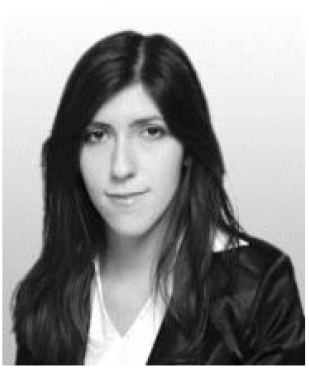

**Ilinka Ivanoska** received the B.Sc. degree from the Faculty of Electrical Engineering and Information Technology, Ss. Cyril and Methodius University, Skopje, North Macedonia, in 2009, and the Ph.D. degree in Computer Science from Faculty of Computer Science and Engineering, Ss. Cyril and Methodius University, Skopje in 2021. She is currently an Associate Professor with the Faculty of Computer Science and Engineering, Ss. Cyril and Methodius University, Skopje. As a professor, she has taught courses in the areas of algorithms and data structures, probability and statistics, object-oriented systems, machine learning, and bio-inspired computing. Her research interests include machine learning and its applications in computational biology, neuroimaging and complex networks.

