## Supplementary Materials for "Task-Dependent Performance of GNNs and Pretrained Models in ADMET Prediction"

Eight message-passing variants (GraphConv, GCN, GAT, GraphSAGE, GIN, TAG, SGC, TransformerConv) were screened on all six endpoints under the main study's scaffold-split pipeline, target transformations, and metrics. Each architecture–endpoint result is the best of five configurations selected on validation (search over hidden dimension, layers, and learning rate), i.e. $8 \times 6 \times 5 = 240$ runs. Noarchitecture dominates: validation ranks span a narrow band (average regression rank 3.0–6.0, Table S3; validation AUC 0.78–0.85 on Tox21 and 0.88–0.92 on hERG,Table S1), every architecture is unstable on the clearance and half-life endpoints, and validation order predicts test performance poorly (the best validation-ranked backbone,GraphSAGE, reaches only mean test $R^2 = 0.10$, while others fall to $-1.57$). GraphConv (validation rank 4.25 of 8, mean test $R^2$ among the lowest) was adopted as a single representative backbone usingvalidation metrics only, so the main-benchmark test evaluation remains unbiased. Table S2 lists the GNN hyperparameter search space,Figs. S1 and S2 the imbalance-aware toxicity diagnostics, and Tables S4 and S5 the HPO-optimizer comparison.

TABLE S1
CLASSIFICATION ARCHITECTURE SCREEN: VALIDATION AUC-ROC AND F1 (BEST OF FIVE VALIDATION-SELECTEDCONFIGURATIONS). BEST AUC IN **BOLD**; TEST DATA NOT USED.

| Architecture | Tox21 (NR-AR) | | hERG | |
|---|---|---|---|---|
| | AUC | F1 | AUC | F1 |
| GraphConv | 0.819 | 0.480 | 0.918 | 0.903 |
| GCN | 0.821 | 0.435 | 0.912 | 0.915 |
| GAT | 0.819 | 0.500 | 0.907 | 0.871 |
| GraphSAGE | **0.854** | 0.421 | 0.903 | 0.918 |
| GIN | 0.844 | 0.500 | 0.884 | 0.933 |
| TAG | 0.821 | 0.444 | 0.892 | 0.897 |
| SGC | 0.829 | 0.500 | **0.920** | 0.875 |
| Transformer | 0.778 | 0.480 | 0.905 | 0.881 |

TABLE S2
GNN HYPERPARAMETER SEARCH SPACE. THE THREE MLP-HEAD WIDTHS ARE SAMPLED INDEPENDENTLY, THENSORTED IN DESCENDING ORDER.

| Hyperparameter | Search Space |
|---|---|
| Hidden dimensions | {64, 96, 128, 192, 256, 384, 512} |
| Number of layers | {3, 4, 5, 6, 7} |
| Learning rate | [1e-4, 1e-2] (log-uniform) |
| Weight decay | [1e-6, 1e-2] (log-uniform) |
| MLP head layer 1 | {128, 192, 256, 384, 512} |
| MLP head layer 2 | {64, 96, 128, 192, 256} |
| MLP head layer 3 | {32, 48, 64, 96, 128} |

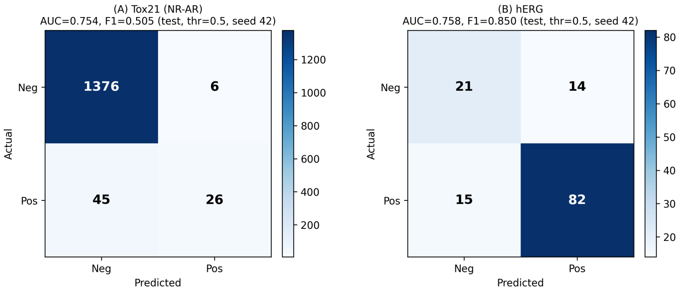


Fig. S1. Confusion matrices for the GNN's validation-selected configuration on the test set(threshold 0.5, seed 42). (A) Tox21 shows majority-class bias under extreme imbalance (F1 = 0.51). (B) hERG is balanced (F1 = 0.85). Across-seed aggregates are in Fig. S2.

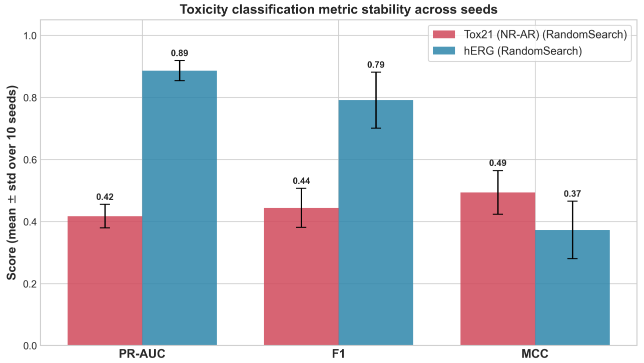


Fig. S2. Across-seed mean ± one standard deviation (ten seeds) of PR-AUC, F1, and MCC.Variability is modest, with F1 and MCC somewhat more variable on hERG than on Tox21.

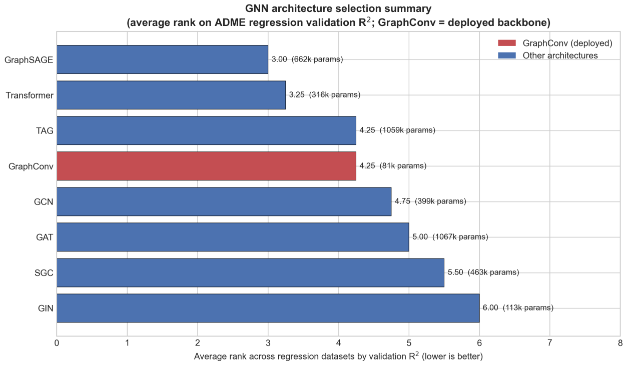


Fig. S3. Average validation-$R^2$ rank across the regression datasets (lower is better); GraphConv (deployed backbone) highlighted. Differences fall within the noise of these small datasets.

TABLE S3

REGRESSION ARCHITECTURE SCREEN: AVERAGE VALIDATION-$R^2$ RANK (LOWER IS BETTER) AND PER-DATASETVALIDATION $R^2$; TEST $R^2$ IS THE MEAN TEST VALUE (POST HOC REFERENCE ONLY, NOT USED FOR SELECTION). GRAPHCONV (DEPLOYED) IN **BOLD**.

| Arch. | Rank | Caco-2 | Half-Life | Hep. | Mic. | Test $R^2$ | Params | Time (s) |
|---|---|---|---|---|---|---|---|---|
| GraphSAGE | 3.00 | 0.597 | 0.434 | 0.316 | 0.175 | 0.096 | 662k | 7.3 |
| Transformer | 3.25 | 0.606 | 0.335 | 0.327 | 0.208 | −0.197 | 316k | 16.4 |
| TAG | 4.25 | 0.582 | 0.407 | 0.228 | 0.279 | 0.226 | 1059k | 12.2 |
| **GraphConv** | 4.25 | 0.575 | 0.460 | 0.267 | 0.081 | −0.673 | 81k | 7.1 |
| GCN | 4.75 | 0.534 | 0.447 | 0.247 | 0.137 | −0.673 | 399k | 12.8 |
| GAT | 5.00 | 0.513 | 0.483 | 0.233 | 0.122 | 0.128 | 1067k | 14.9 |
| SGC | 5.50 | 0.560 | 0.426 | 0.194 | 0.159 | 0.006 | 463k | 12.6 |
| GIN | 6.00 | 0.625 | 0.368 | 0.157 | 0.058 | −1.572 | 113k | 6.2 |

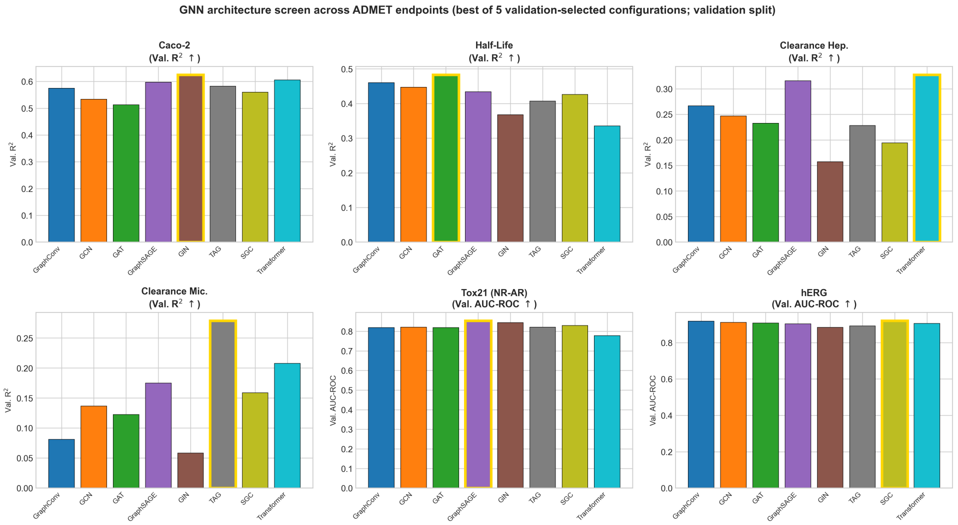


Fig. S4. Per-dataset architecture comparison (validation AUC-ROC for classification, $R^2$ for regression). Clearance and half-life endpoints are unstable across all eight architectures;structure-driven endpoints cluster narrowly.

TABLE S4

HPO STRATEGIES VS. RANDOM SEARCH: EACH OPTIMIZER'S VALIDATION-SELECTED CONFIGURATION EVALUATED ONCE ON TEST ACROSS $n = 6$ DATASETS. **W/T/L**: WINS/TIES/LOSSES, WHERE A PAIRED DIFFERENCE SMALLERTHAN $10^{-3}$ IN ABSOLUTE VALUE IS COUNTED AS A TIE; **SIGN** $p$: EXACT TWO-SIDED SIGN TEST (TIES DROPPED); **NORM. EFFECT**: PER-ENDPOINT DIFFERENCE DIVIDED BY THE SEVEN-OPTIMIZER SPREAD,AVERAGED; **CLIP/FLOOR-0**: MEAN PAIRED DIFFERENCE WITH $R^2$ CLIPPED TO $[-1, 1]$ / FLOORED AT 0. 95% CIS ARE 10,000-RESAMPLE PERCENTILE BOOTSTRAPS. NO OPTIMIZER IS SIGNIFICANT UNDER THE SIGNTEST AND EVERY SCALE-FREE/CLIPPED INTERVAL SPANS ZERO; ONLY THE FLOOR-0 VARIANT SHOWS MARGINAL EFFECTS (ABC SLIGHTLY +, SA SLIGHTLY −), NEITHER INDICATING A CONSISTENT WINNER.

| Optimizer | W/T/L | Sign $p$ | Norm. effect [95% CI] | Clip $[-1, 1]$ mean [95% CI] | Floor-0 mean [95% CI] |
|---|---|---|---|---|---|
| PSO | 2/0/4 | 0.69 | −0.16 [−1.35, +1.25] | +0.174 [−0.084, +0.612] | +0.033 [−0.036, +0.124] |
| ABC | 4/1/1 | 0.375 | +0.52 [−0.45, +1.38] | +0.026 [−0.292, +0.311] | +0.032 [+0.000, +0.091] |
| GA | 3/0/3 | 1.00 | +0.01 [−1.41, +1.40] | +0.169 [−0.067, +0.536] | +0.020 [−0.040, +0.086] |
| SA | 1/0/5 | 0.22 | −0.77 [−1.82, +0.55] | −0.135 [−0.471, +0.171] | −0.018 [−0.031, −0.006] |
| HC | 3/0/3 | 1.00 | +0.02 [−1.13, +1.37] | +0.034 [−0.440, +0.556] | +0.014 [−0.021, +0.062] |
| TPE | 2/0/4 | 0.69 | +0.01 [−1.02, +1.16] | +0.105 [−0.075, +0.329] | +0.027 [−0.024, +0.114] |

TABLE S5

TEST PERFORMANCE OF EACH OPTIMIZER'S VALIDATION-SELECTED CONFIGURATION (REGRESSION $R^2$; CLASSIFICATION AUC-ROC), SHOWN FOR TRANSPARENCY; STATISTICS VS. RANDOM SEARCH ARE INTABLE S4. CLEARANCE $R^2$ IS HIGHLY UNSTABLE, SO PER-OPTIMIZER REGRESSION VALUES SHOULD NOT BE OVER-INTERPRETED.

| Dataset | PSO | ABC | GA | SA | HC | Random | TPE |
|---|---|---|---|---|---|---|---|
| Caco-2 ($R^2$) | 0.508 | 0.553 | 0.450 | 0.519 | 0.519 | 0.552 | 0.515 |
| Half-Life ($R^2$) | 0.119 | 0.211 | 0.193 | −0.285 | 0.037 | 0.035 | 0.274 |
| Hepatocyte ($R^2$) | −0.269 | −0.762 | −0.219 | −1.351 | −0.989 | −0.114 | −0.283 |
| Microsomal ($R^2$) | 0.235 | −0.387 | 0.058 | −0.529 | 0.130 | −3.288 | −0.365 |
| Tox21 (AUC) | 0.739 | 0.757 | 0.730 | 0.739 | 0.758 | 0.743 | 0.726 |
| hERG (AUC) | 0.738 | 0.810 | 0.830 | 0.771 | 0.777 | 0.810 | 0.787 |
